# Cognitive Improvements after Intermittent Deep Brain Stimulation of the Nucleus Basalis of Meynert in a Transgenic Rat Model for Alzheimer’s disease; a Preliminary Approach

**DOI:** 10.1101/600296

**Authors:** Philippos Koulousakis, Daniel van den Hove, Veerle Visser-Vandewalle, Thibaut Sesia

## Abstract

**Background:** Deep brain stimulation (DBS) of the nucleus basalis of Meynert (NBM) has shown to have promising results in a pilot study with patients suffering from Alzheimer’s disease (AD). A recent study in aged monkeys shows a novel intermittent stimulation pattern to have superior cognitive benefits over continuous paradigms.

**Objective/Hypothesis:** We aimed at comparing the cognitive effects elicited by intermittent and continuous NBM stimulation paradigms in an animal model for AD (TgF344-AD rat line; TG), i.e. rats expressing mutant human amyloid precursor protein (APPsw) and presenilin 1 (PS1ΔE9) genes, each independent causes of early-onset familial AD.

**Methods:** In this exploratory study, aged APP/PS1 rats were tested pre-, and post implantation with several stimulation parameters, i.e. unilateral- or bilateral-intermittent, and bilateral-continuous, while performing various behavioral tasks (open field, novel object recognition, and modified Barnes maze).

**Results and Conclusion:** Bilateral-intermittent NBM DBS allowed aged TG rats to perform better and maintain their performance longer in a spatial memory task, as compared to other conditions. These data support the notion that NBM DBS could be further refined in the clinic, thereby improving patient care.

## Introduction

Continuous deep brain stimulation (DBS) of the nucleus basalis of Meynert (NBM) has been shown to be safe and exert pro-cognitive effects in patients diagnosed with Alzheimer’s disease (1–5). The stimulation is likely to drive the dense population of cholinergic neurons of the NBM, functionally associated with control of attention and maintenance of arousal, both key functions for learning and memory formation (6–11). Studies in aged rhesus monkeys show *intermittent* NBM stimulation to induce supranormal working memory and sustained attention (12,13), suggesting that the current pattern of stimulation used in the clinic may not be optimal (14). Accordingly, in the present study, we aimed at comparing the cognitive benefits elicited by intermittent and continuous NBM stimulation in an animal model for AD (TgF344-AD rat line; TG), i.e. rats expressing mutant human amyloid precursor protein (APPsw) and presenilin 1 (PS1ΔE9) genes, both causal to early-onset familial AD in humans (15,16). This exploratory study aims at testing the potential of intermittent stimulation for patients suffering from AD in a diseased stage. Therefore, transgenic rats matching the age of clinical diagnosis will be stimulated with various stimulation parameters and performances compared to pre-surgery baseline (17).

## Materials and Methods

### Animals

Twelve 18-month old TG male rats were housed in a normal day-night cycle with *ad libitum* access to water and food. Experimental schedule lasted for three months, with each animal undergoing the same stimulation paradigm at the same time. Based on the data available on this rat line, the age that model the most time during which a patient is most like to be diagnosed with AD is between 16-20 months (17–19). This study was performed in accordance with German regulations and legal requirements and was approved by the local government authorities (Landesamt für Natur, Umwelt und Verbraucherschutz Nordrhein-Westfalen (LANUV NRW), North Rhine-Westphalia, Germany).

**Fig 1.**
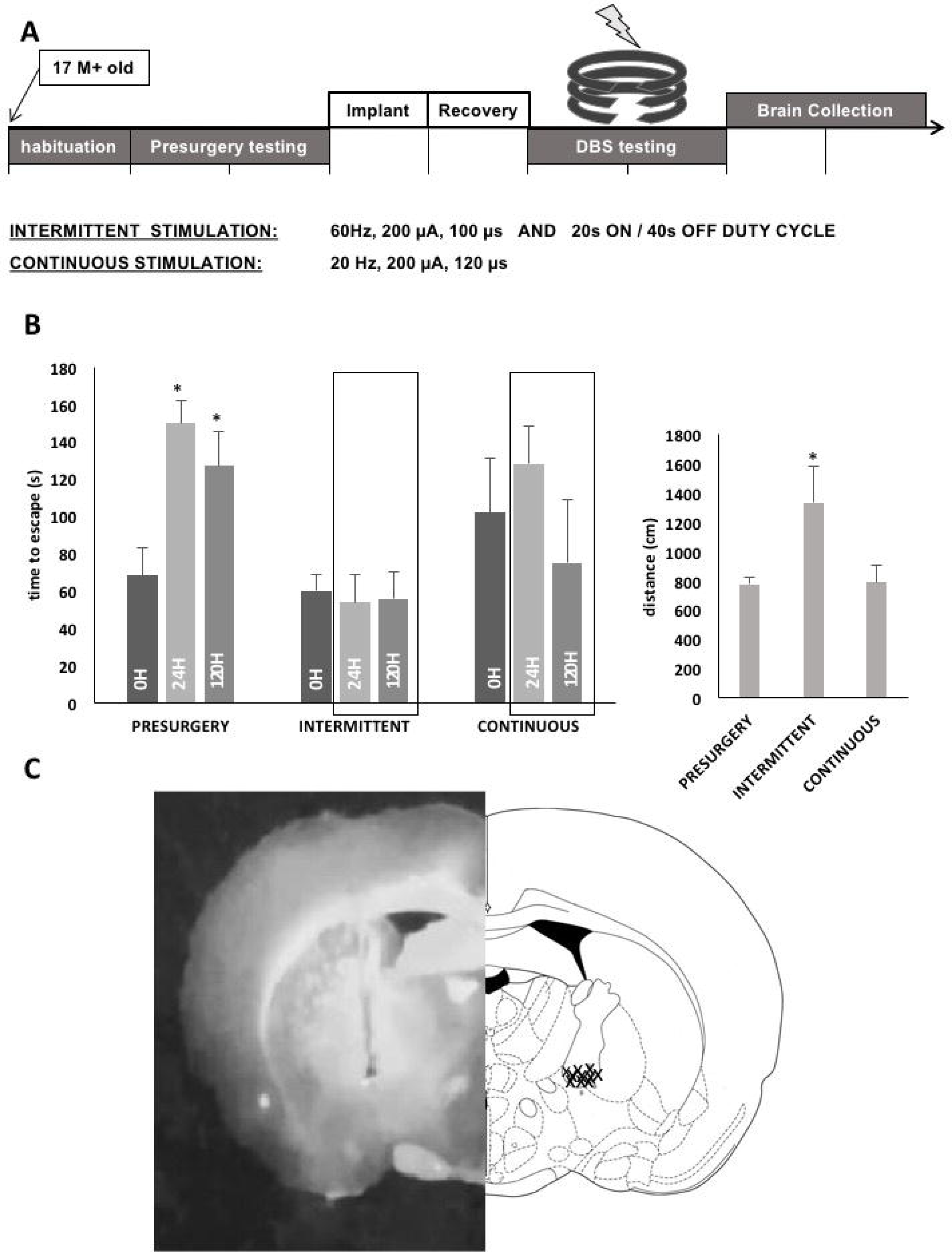
Experimental schedule outline shown regarding the experimental schedule from habituation to brain collection. (A) Effect of intermittent NBM DBS on spatial memory and locomotion. Time to escape (left) and distance covered (right) of aged male Tgf344-AD rats performing a MBM test pre-surgery, with intermittent or continuous stimulation (training day TD3; test 1 1D after training; test 2 5D after training), or open field testing. Each animal performs all conditions. A difference from the respective training condition is indicated by asterisks (MBM Presurgery ranked-sum test latency: Z=11;3 n=10 P <.05* two-tailed; errors: Z=5;3 n=10 P <.05*; OF Presurgery ranked-sum test OF: distance: Z=3,6; n=10; P<.05). (B) Electrode localization. Right side schematic representation. Left side fresh section showing electrode tract. (C)

### Surgery and postoperative care

Rats were injected with Carprofen (5,0mg/kg) and Buprenophrin (0,05mg/kg) 60 minutes presurgery. Rats were anaesthetised by gaseous isoflurane before bilateral implantation of the stimulating electrodes into the NBM (coordinates in mm from bregma: AP= −1,44; ML= +/-2,88; DV= 7,4(16). Electrodes were implanted at a 6° angle and 4 micro screws were used to stabilize the implant. Carprofen (5,0mg/kg/day) was administered for 2 days post-surgery. Rats recovered for at least a week before the first stimulation.

### Stimulation Paradigms

Animals were stimulated intermittently (uni- or bilaterally) or continuously (bilaterally only) using bipolar platinum-iridium stimulating electrodes (Plastics One, Roanoke, VA, USA). Positive monopolar pulses were generated by a Master-8 stimulator paired to 2 -Flex stimulus isolators (A.M.P.I., Jerusalem, Israel). The pulse width was set at 100μs, the intensity at 200μA and the frequency at 60Hz for intermittent stimulation, and 20Hz for continuous stimulation. The duty cycle of intermittent stimulation pattern was 20 seconds ON and 40 seconds OFF.

### Behavioral Testing

Twelve animals were tested in an open field test (OF), a novel object recognition task (NOR), and a modified Barnes maze task (MBM) (5 days; 3 training/ 2 probe trials), pre- and post-implantation. Behavioral testing was performed between 8 AM and 2 PM. Animals were brought into the testing room 30 minutes beforehand. Lighting was set at 80 lumen. Mazes were cleaned before and after each trial with 10% white vinegar solution. The order in which animals were tested was randomized daily. Post-implantation, rats were stimulated during testing. One animal fell off the maze during MBM training. The implant was damaged, and the animal was removed from the study.

### Modified Barnes Maze Task

In this spatial memory test, rats were to escape a stressful environment (bright light) into an escape box. The maze consisted of a circular dark-grey platform (120 cm diameter) with 20 holes evenly placed around the perimeter; only one hole led to the escape box. The escape box location remained consistent throughout the trial (15). Rats were trained twice per day for three days, then tested one and five days later (probe trials). The time to reach the escape box and the number of errors, explorations of holes other than the escape hole, was recorded.

### Open Field Test

The arena used consisted of a 60 cm high −100 cm square box. Rats were placed into the centre of the maze and were allowed to explore the arena for 10 minutes. Total distance moved, and total amount of time spent in centre vs border zones was measured.

### Novel Object Recognition Task/Test

In the same arena, 2 identical objects were placed 25cm off the walls. Animals were placed in the maze facing away from objects and allowed to explore for 3 minutes. Afterwards, rats were returned to their home cage for 20 minutes. Object sets were permuted with a novel object and a clean familiar object. Rats were returned to the maze to explore for 3 minutes. Total amount of time spent exploring each object was measured for both trials, as well as discrimination scores (d1, d2)(18).

### Statistical Analysis

Data were compared with a Wilcoxon signed-ranked test. Individual performances of animals were ranked and rankings within conditions compared. For MBM analysis the difference in rankings between the final day of training and the probe trials after 1 and 5 days respectively were compared.

## Results

Rats spent significantly more time in border zones than the center zone of the OF throughout all conditions without significant differences between conditions (Z=0 p<.05; SEM **presurgery:** 7.8; **unilateral:** 2.79; **bilateral:** 2.82; **continuous:** 1.25). Rats covered more distance in the OF under unilateral (Z=3 p<.05; μ=1319.125, SEM = 48.68) and bilateral (Z=3 p<.05; μ= 1335.488, SEM = 247.45) intermittent, but not continuous stimulation (Z=6 p>.05; μ= 791.9, SEM = 110.468), when compared to presurgery (μ= 775.67, SEM = 48.6).

NOR results showed low exploration throughout conditions (e1= total exploration 1^st^ trial; e2= total exploration 2^nd^ trial: **presurgery:** e1=5.09; e2=8.9; **unilateral:** e1=3.2; e2=3.6; **bilateral:** e1=3.75; e2=4; **continuous:** e1=2.5; e2=3.0). Discrimination index d2 shows no significant novel object recognition for any condition when compared against zero **(presurgery:** d2 = 0.04, p<.05; **bilateral:** d2 = 0.35, p<.05 **unilateral:** d2 = 0, p<.05; **continuous:** d2:0.11, p<.05).

Rats made significantly more errors presurgery, 1 day (Z=3; p<.05), as well as 5 days (Z=11; p<.05) after the final training in the MBM when compared to the final day of training (TD3: μ= 8.22, SEM = 0.76; T1D: μ= 19.727, SEM = 3.0; T5D: μ*=* 15.72, SEM = 3.5). There was no significant difference in errors made comparing final training day with 1 day or 5 days after training throughout intermittent stimulation conditions **(unilateral:** Z=18;17; p>.05; TD3: μ= 8.18,SEM = 1.7; T1D: μ = 9.375,SEM = 2.8; T5D: μ = 9, SEM = 2.25; **bilateral:** Z=28;28; p>.05; TD3: μ = 9.83, SEM = 1.8; T1D: μ = 10.90, SEM = 1.7; T5D: μ = 8.18, SEM = 1.8) and continuous stimulation (Z=2;1 p<.05; TD3: μ = 10.125,SEM = 2.98; T1D: μ = 11;SEM = 0.70; T5D: μ = 4). However, animal immobility was observed during continuous stimulation.

Rats needed significantly longer to escape the maze presurgery, 1 day (Z=2; p<.05), as well as 5 days (Z=0; p<.05) after the final training day when compared to the final day of training (TD3: μ =68.2, SEM = 15; T1D: μ = 149.9, SEM = 12.14; T5D: μ = 126.9, SEM = 18.9). When stimulated intermittently unilaterally, rats maintained their performance 1 day (Z=15; p>.05), and 5 days (Z=5; p>.05) after the final day of training (TD3: μ = 99.75, SEM = 22.43; T1D: μ = 81.75, SEM = 19.93; T5D: μ = 42.125, SEM = 13.1). Similarly, rats stimulated bilaterally intermittently maintained performance 1 day (Z=16; p>.05) as well 5 days (Z=15.5; p>.05) after training (TD3: μ = 60, SEM = 8.8; T1D: μ = 53.89, SEM = 14.61; T5D: μ = 55.56, SEM = 14.78). Rats stimulated continuously did not maintain their performance 1 day (Z=0; p<.05) or 5 days (Z=2; p<.05) after training (TD3: μ = 102 SEM = 28.9; T1D: μ = 128, SEM = 20; T5D: μ = 75, SEM = 34.03).

## Discussion

In this experiment we tested the hypothesis that bilateral intermittent stimulation of the NBM leads to superior cognitive benefits in aged TG rats when compared to continuous stimulation. Data indicate that locomotion was increased during unilateral, as well as bilateral intermittent stimulation, suggested by distance covered in an open field, when compared to presurgery. Continuous stimulation did not lead to an increased distance covered when compared to presurgery. Moreover, anxiety levels in the OF were unaffected by stimulation patterns when compared to presurgery. NOR data indicate a low overall exploration of the objects indicated by e1 and e2 scores. Novel object recognition memory was not achieved at any condition.

Data suggest that spatial memory can be improved with bilateral intermittent NBM-DBS. Partial stabilization was also achieved with unilateral intermittent stimulation. Continuous stimulation did not lead to statistical improvement of performance, possibly due to the prevalent immobility observed during continuous stimulation.

These data are promising; additionally, limitations need to be considered. Of special note is the age of the animals. The overall experimental schedule lasted 3 months: the sample size decreased over time. While 11 animals were available for bilateral intermittent stimulation, 4 remained for continuous stimulation. Two animals had to be excluded due to health issues. Five animals developed a kindling effect upon continuous stimulation. Further research including more animals is needed to evaluate if this is e.g. a direct effect of the stimulation pattern or a carry-over effect resulting from prolonged stimulation. This side effect has never been reported in patients though.

## Conclusion

Bilateral intermittent NBM DBS allows aged TG rats to perform better and maintain their performance longer in a spatial memory task when compared to presurgery. These data support that the NBM DBS technique can be further refined in the clinic, thereby improving patient care.

## Acknowledgment

This work was supported by the Department of Stereotactic and Functional Neurosurgery of the University Hospital of Cologne.

